# Shrinkage estimation of gene interaction networks in single-cell RNA sequencing data

**DOI:** 10.1101/2024.03.20.585951

**Authors:** Duong H.T. Vo, Thomas Thorne

## Abstract

Gene interaction networks are graphs in which nodes represent genes and edges represent functional interactions between them. These interactions can be at multiple levels, for instance, gene regulation, protein-protein interaction, or metabolic pathways. To analyse gene interaction networks at a large scale, gene co-expression network analysis is often applied on high-throughput gene expression data such as RNA sequencing data. With the advance in sequencing technology, expression of genes can be measured in individual cells. Single-cell RNA sequencing (scRNAseq) provides insights of cellular development, differentiation and characteristics at transcriptomic level. High sparsity and high-dimensional data structure pose challenges in scRNAseq data analysis. In this study, a sparse inverse covariance matrix estimation framework for scRNAseq data is developed to capture direct functional interactions between genes. Comparative analyses highlight high performance and fast computation of Stein-type shrinkage in high-dimensional data using simulated scRNAseq data. Data transformation approaches also show improvement in performance of shrinkage methods in non-Gaussian distributed data. Zero-inflated modelling of scRNAseq data based on a negative binomial distribution enhances shrinkage performance in zero-inflated data without interference on non zeroinflated count data. The optimal zero-inflated Stein-type shrinkage framework is applied on experimental scRNAseq data which demonstrates its potential to construct sparser gene interaction networks with higher precision.

**Availability and implementation:** https://github.com/calathea24/ZINBGraphicalModel

## 1 Introduction

In a biological system, specific cellular activities or functions are often carried out by interactions between genes and their products. These interactions form dynamic, complex networks which can result in changes in higher-level biological systems such as tissues and organs. Constructing and characterising these gene interaction networks are of paramount importance for guiding experimental designs, biomarker identification in diagnostic and prognostic research, target in drug discovery and development, and understanding the biological processes in an organism [1].

In gene network analysis, nodes represent genes or gene products and edges represent pair-wise relationship between them. Data from high-throughput experiments such as RNA sequencing (RNAseq) are often used to construct global gene co-expression networks which might provide insights of gene interaction networks [2]. RNA sequencing uses nextgeneration sequencing to quantify the amount of RNA molecules in a biological sample and reveal difference in gene expression between different samples [3]. The expression value of each gene in RNAseq data represents average gene expression of a large population of cells [4]. With advances in sequencing technologies, gene expression data are able to be extracted from individual cells in specific cell types [5]. The power and potential of single-cell RNA sequencing (scRNAseq) have been demonstrated in cell development and differentiation research [6, 7]. Computational methods to harness this potential are still in their infancy as methods developed for bulk RNA-seq data are often applied [3].

Differences in properties of scRNAseq data from bulk RNAseq data have been highlighted, including the potential for dropout events which introduce technical noise in the data, stronger overdispersion between individual cells, heterogeneity of sequenced cells due to presence of different cell populations, and different cell states and cell cycles [4]. These differences can hinder performance of existing computational tools for sequencing data analysis [3]. Therefore, computational methods specific for scRNAseq data with high performance are essential to facilitate applications of single-cell RNA sequencing in research.

In scRNAseq data analysis, the term dropout is used to describe the prevalence of excessive zero counts [8]. Dropout events are defined as the situation when expression of a gene is detected in some cells but absent in other cells of the same cell types in the same experiment [9]. Multiple factors are suggested to cause dropout events such as failure in capturing and amplifying mRNAs or limitation in sequencing depth [10]. For example, in a quantitative assessment of scRNAseq methods, a low-expressed gene TERT was only detected by quantitative polymerase chain reaction (qPCR) compared to scRNAseq [5]. Stochastic gene expression or existence of a new underlying cell subpopulation are also potential dropout-event factors [11, 12]. In the current literature, there are debates about whether dropout events occur in scRNAseq protocols using unique molecular identifiers (UMI) or whether the usage of zero-inflated modelling in scRNAseq data analysis improves performance [10, 13].

Covariance or correlation matrices from Pearson or Spearman correlation tests are used widely to generate connectivity graphs of gene co-expression. One main limitation is correlation analysis not only captures pairwise correlation between genes but also associations contributed by third-party or global effects [14]. To overcome this limit, graphical models can be applied to measure the direct interaction between genes or variables. A class of these, called Gaussian Graphical Models, make use of the partial correlation matrix, which can be calculated from the inverse covariance (precision) matrix. However, when the number of samples is comparable or smaller than the number of variables, the sample covariance matrix inherits a lot of estimation errors [15]. To solve this issue, shrinkage approaches can be adopted in which the precision matrix is assumed to be sparse.

In this study, we compare the performance and computation time of Stein-type shrinkage methods to Lasso-type shrinkage methods using simulated scRNAseq data. We build on the existing Stein-type shrinkage approach of Schäfer and Strimmer [16] by integrating zero-inflated negative binomial mixture modelling, and identifying the optimal data-transformation scheme for scRNAseq counts. Finally, our suggested workflow of zeroinflated Stein-type shrinkage is applied in experimental scRNAseq data of *Saccharomyces cerevisiae* and *Schizosaccharomyces pombe*.

## 2 Methods

Graphical models are graphs in which nodes are random variables and the presence of edges represent conditional dependence or “direct links” between them [17]. Two random variables, X and Y, are considered to be independent conditionally on variable Z if their conditional probability meets:

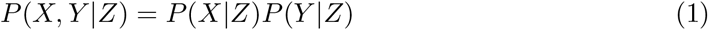

Gaussian Graphical Models (GGMs) are a popular form of undirected graphical model in which a random variable X follows a multivariate Gaussian distribution *𝒩* (0, Σ). The covariance matrix Σ is unknown and nonsingular [17].

A GGM is represented by a partial correlation matrix P, as zero values in P correspond to conditional independence between variables [18]. Partial correlation coefficients measure linear relationships between pairs of random variables (*i* and *j*) which corrects for the effect from other variables (conditional dependence) [19]. The partial correlation coefficient *ρ*_*ij*_ can be calculated based on Ω, the inverse of covariance matrix Σ:

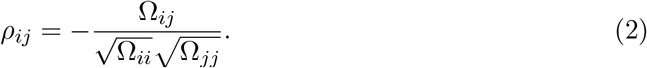

In practice, the most common covariance estimator is the sample covariance matrix. However, when the number of features (p) exceeds the number of samples (n), the sample covariance matrix becomes singular and non-invertible [20]. In this situation, shrinkage techniques are often adopted to generate an invertible estimated covariance matrix for calculating the partial correlation matrix.

There are two common approaches for estimation of sparse partial correlation matrices, using *L*_1_/lasso regularization in a maximum likelihood framework (Lasso-type) and linear shrinkage to shrink the sample covariance matrix towards a structured target matrix (Stein-type). In Lasso-type shrinkage, let *S* be the sample covariance matrix, and Θ be the estimated inverse covariance matrix, then the penalized log-likelihood function to maximize is [21]:

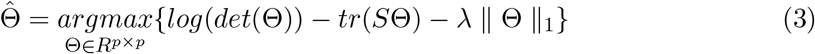

where *λ* is the amount of shrinkage applied and chosen by users in implementation. On the other hand, Stein-type shrinkage comprises three components, an empirical covariance matrix *S*, a shrinkage target matrix *T* and a shrinkage constant *α*. The estimated covariance matrix is computed through a linear combination [15]:

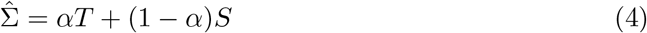

In this study, Lasso-type shrinkage is implemented with the graphical lasso (glasso) [22] and Meinshausen-Buhlmann (mb) [23] algorithms, using the huge R package [24]. Stein-type shrinkage is applied using identity matrix as target matrix and shrinkage intensity (*α*) is calculated based on results of [16] and [25] studies which are named as GeneNet and Fisher2011 algorithms, respectively (Supplementary materials 1-3).

### 2.1 Data transformation and p-value calculation for Stein-type shrinkage workflows

In Gaussian graphical models, normality is a standard assumption about distribution of count data [26]. However, RNAseq data has specific properties such as skewness, extreme values, mean-variance dependency which deviates from the normal distribution assumption [27, 28]. To improve the performance of shrinkage methods, data transformation approaches are recommended [27].

Three data transformation methods are compared in terms of enhancing performance of Stein-type shrinkage. We consider log transformation as log(*count* + 1), which is highly used in biological data analysis to convert skew distribution towards normally distributed data [27]. We also apply a semiparametric Gaussian copula or nonparanormal transformation, which was developed for high-dimensional graphical modelling by using a Gaussian copula with Winsorized truncation [29]. This algorithm is implemented using huge.npn function from huge R package [24]. Lastly, we also consider the empirical copula Φ^*–*1^(*ECDF* (*count*)) which is similar to the nonparanormal approach and includes a Gaussian copula based on the empirical cumulative distribution of the data without truncation.

Compared to Lasso-type shrinkage methods, Stein-type shrinkage returns an estimated partial correlation matrix which requires significance tests to generate a final graph. Generally, partial correlation coefficients are treated as Pearson’s correlation coefficients and tested using t-statistics [30, 31]. This has been applied in the GeneNet R package for Stein-type shrinkage. To take into account the effect of shrinkage and its intensity on partial correlation matrix, Bernal V. et al. proposed two new probability densities for significance test which are referred to as shrunkv1 and shrunkv2 in this study [19, 31].

### 2.2 Modifications in shrinkage frameworks for zero-inflated counts

To fully exploit the potential of scRNAseq data, one main challenge to overcome is the presence of excessive zeros in count data due to dropout events. Dropout events in scRNAseq data are defined as the situation where the expression of a gene is detected in some cells but absent in other cells of the same cell type [9]. Therefore, to reduce estimation errors in the sample covariance matrix and stratify biological zeros from dropout zero counts, we propose to use zero-inflated z score calculation based on zero-inflated negative binomial (ZINB) modelling of the count data in shrinkage framework. Details of the process is illustrated in Supplementary material 4.

In the first step, a ZINB model is fitted for expression counts of each gene *j* in cell *i* (*Y*_*ij*_) by the L-BFGS-B algorithm using the `optim` R function to derive maximum likelihood estimates of parameters of the model:

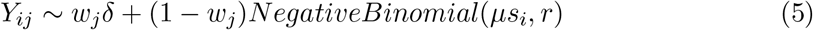

where *μ* is mean, s is vector of constant scaling factors (see Supplementary material 4) for each cell and *r* is the size parameter of the negative binomial distribution. Parameter *w*_*j*_ balances negative binomial model accounting for true expression counts and delta function. For count of gene j in cell i, delta function returns 0 value for all non-zero counts.

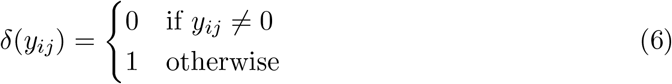

Next, a non-detection rate (*d*_*ij*_) is calculated based on the optimized ZINB model of each gene. For each count of gene j in cell i:

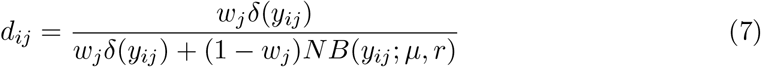

To select zero counts in the data resulting from dropout events, a threshold *t* with default value 0.5 is applied. After zero counts potentially resulting from dropout events are identified, a nonparanormal transformation is applied to the count data [29].

Zero-inflated z-scores are then calculated in which z-scores of zero counts identified as dropout events are set to 0. To calculate zero-inflated z-scores of nonparanormal transformed data X of gene j in cell i we use:

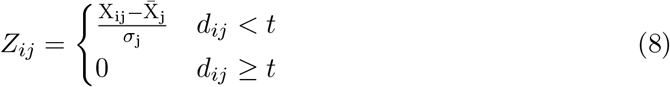

Our proposed methodology, referred to below as ZIGeneNet, fits the ZINB mixture model given above to each gene, and then uses nonparanormal transformation of the data and the derived non-detection rates to calculate the zero-inflated z-scores *Z*_*ij*_ of each count which are then used in empirical covariance matrix *S* calculation of a Stein-type shrinkage approach. The shrinkage coefficient *α* is chosen using the methodology of [16].

Significant edges are detected using correlation test statistics from fdrtool R package [30]. The workflow is outlined in Supplementary Figure S5A.

## 3 Results and Discussion

Simulated data for scRNAseq was generated using an algorithm based on the ESCO simulation tool [32]. A simplified simulation version with incorporated partial correlation matrix is illustrated in Supplementary Figure S6. The simplified scRNAseq model is for homogeneous cell populations when there are no outlier cells and all cells belong to one cell group. Genes in the gene network are simulated to be expressed in at least 60% of cells. This matches guidelines in experimental scRNAseq data analysis in which genes not expressed in a certain number of cells are excluded for downstream analysis [33]. To simulate regulatory interactions between genes, a correlation matrix is calculated based on a partial correlation matrix provided as an input.

Simulation of zero inflation in ESCO is based on mean of gene expression in which genes with higher expression mean are less likely to have zero-inflated counts [32]. This is consistent with current reports about relation between dropout event and level of gene expression [9].

### 3.1 Data transformation and p-value estimation approaches in Stein-type shrinkage

Firstly, different methods of data transformation (pre-shrinkage step) and p-value calculation (post-shrinkage step) are explored to construct an optimal Stein-type shrinkage workflow which can be used to implement in gene network analysis of scRNAseq data. Results of the comparison between three transformation methods we consider, versus no transformation in a Stein-type shrinkage workflow when the number of genes and cells are equal to 200 and sequencing depth is 50000 reads in scRNAseq data simulation are highlighted in Supplementary Figure S7A & S7B. The nonparanormal approach outperforms and increases performance of Fisher2011 from MCC *≃* 0.346 to MCC *≃* 0.681 while MCC value is improved from 0.642 to 0.683 in GeneNet case. The strong assumption of normal distribution in Fisher2011 is re-highlighted by the larger difference in performance of Fisher2011 with and without data transformation compared to GeneNet.

Secondly, performance of shrunkv1 and shrunkv2 p-value tests are compared against a significance test using t-statistics and Monte Carlo p-value calculation in scRNAseq simulated data with 200 genes. Ranging from n=60 to n=200, the t-statistics method performs best in all tested methods using GeneNet shrinkage algorithm (Supplementary Figure S7C). Therefore, nonparanormal and t-statistics approaches are used in data transformation and p-value calculation steps of the optimised Stein-type shrinkage workflow.

### 3.2 Performance comparison of shrinkage methods on nonzero-inflated data

Performance of shrinkage algorithms are then analysed on simulated scRNAseq data, applying a nonparanormal transformation. Matthews correlation cofficient (MCC) is used for evaluation. Different sequencing depth (25000, 50000, 500000 reads) are simulated and performance results are summarized in Figure 1. We find that for low numbers of samples (n = 60 to 200) and high sequencing depth (500 000 reads), Stein-type algorithms have better performance than Lasso-type algorithms. Noticeably, Fisher2011 has slightly lower performance than GeneNet in case of 25 000 and 50 000 depths.

**Figure 1:**
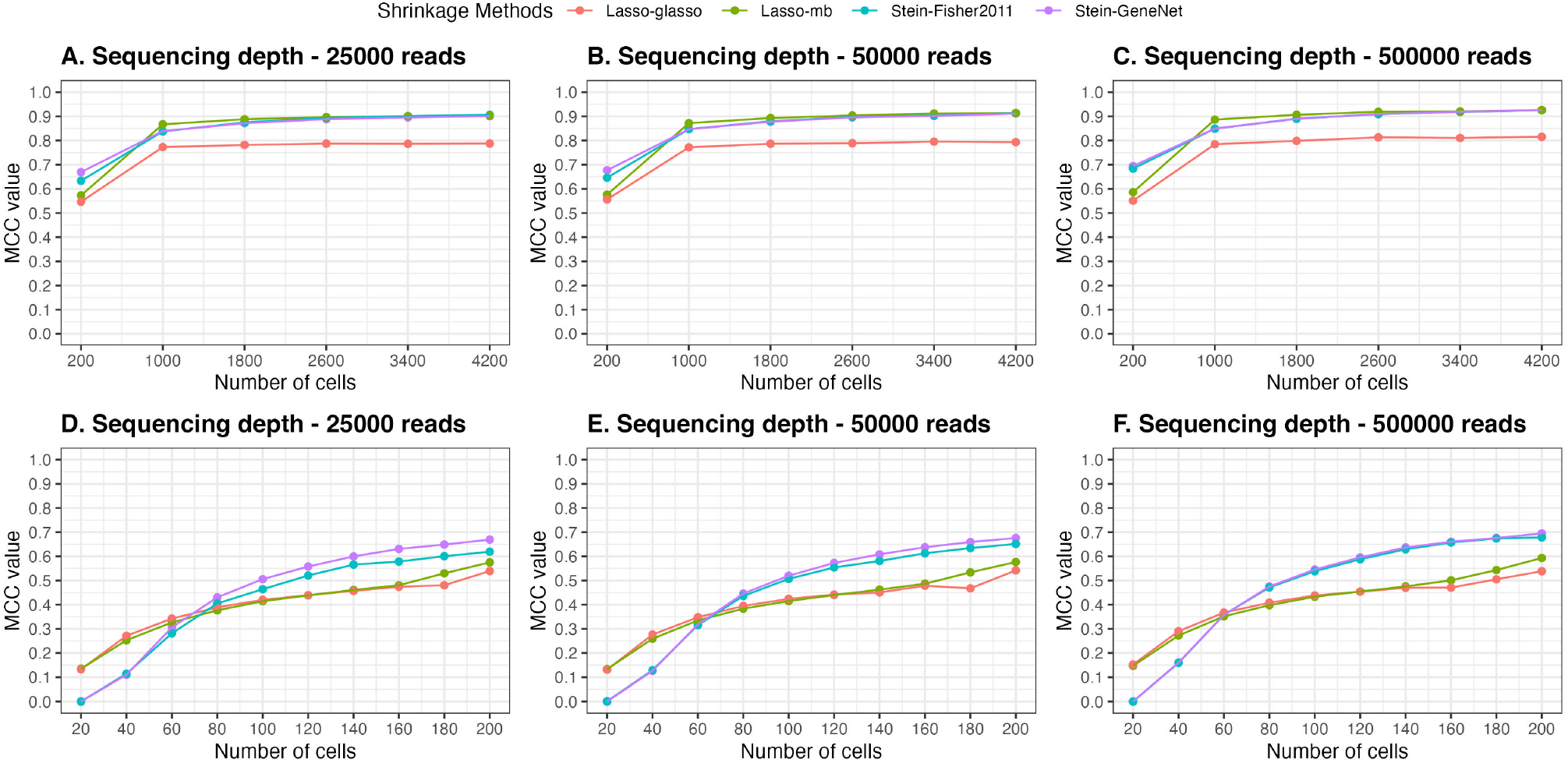
Performance of Stein-type versus Lasso-type shrinkage estimators in scRNAseq simulated data. Different sequencing depths (25000, 50000, 500000 reads) were simulated in the study (iterations: 50).

When designing single-cell RNA sequencing experiments, two main considerations are the number of cells and sequencing depth to sequence individual cells [34]. Increasing sequencing depth can potentially provide more information of gene transcription and reduce technical noise [35]. Interestingly, in simulation, when the number of cells is above 1000, performance of all methods approaches plateau, for example, MCC *≥* 0.8 at sequencing depth of 500 000 reads. In this study, to broaden the application of graphical model, high-dimensional data with “small n large p” are main focus.

Regarding computational cost, computing time of each algorithms is measured using workflows from Supplementary Figure S3 & S4 without a data transformation step on normally distributed data. The simulated data is constructed so that the number of cells is half of the number of genes. Results of average computational time after 10 iterations are shown in Table 1. These results highlight that Stein-type shrinkage methods can perform over 1000 times faster the Lasso-type approaches in some cases.

**Table 1:** Computational time between Stein-type and Lasso-type shrinkage algorithms (seconds). All analyses were performed on Macbook Pro computer with M2 chip, 8GB of RAM and MacOS operating system (10 iterations, p number of features/genes, n number of observations/cells).

| $p$ | $n$ | GeneNet | Fisher2011 | mb | glasso |
| --- | --- | --- | --- | --- | --- |
| 100 | 50 | 0.046 | 0.050 | 41.716 | 52.312 |
| 200 | 100 | 0.148 | 0.144 | 90.048 | 147.519 |
| 400 | 200 | 0.610 | 0.668 | 413.801 | 849.654 |

### 3.3 Zero-inflated approach for Stein-type sparse inverse covariance estimation

Presence of zero counts in experimental scRNAseq data of cells from the same cell type is highlighted in Supplementary Figure S8 and S9 in which gene mean is calculated based on observed/non-zero counts. The plots illustrate that genes with the same gene mean have different proportion of zero counts. This suggests that there might be a subset of genes being affected by dropouts during single-cell experiments.

To consider the impact of modelling zero-inflation of counts for genes that are not affected by dropout, performance of the zero-inflated Stein-type shrinkage framework is tested in simulated scRNAseq data with and without zero inflation (Supplementary Figure S10 & S11). Using GeneNet and Fisher2011 with 200 genes and 100 cells, there is no significant difference in performance of ZI-integrated and non-ZI shrinkage frameworks (p-values of t-test: 0.072 and 0.443, respectively).

The zero-inflated integrated Stein-type shrinkage framework is also compared to an existing graphical model method for sparse gene co-expression network analysis of scRNAseq, scLink [36]. In summary, scLink transforms count data, and uses a Gamma-Normal mixture distribution model to detect missing-value zero counts. Pearson correlation coefficients are calculated based on high-confidence counts and used as input for Lasso-type shrinkage. The Bayesian Information Criterion (BIC) is recommended to choose the value of the regularization parameters [36]. We apply scLink using the built-in functions for data transformation and network estimation (scLink R package), and the optimal graph is chosen by selecting the smallest BIC value. Compared to our Stein-type shrinkage approach, scLink differs mainly in the zero-inflated modelling step (Gamma-Normal versus Negative binomial mixture model), and sparse inverse covariance matrix estimation step (Lasso-type versus Stein-type shrinkage estimator). Performance of scLink versus the proposed ZI shrinkage model is shown in Figure 2 using ZI simulated scRNAseq data. In all cases (n=50, 100, 400 with p=200), ZIGeneNet or the mb shrinkage framework taking into account zero-inflation outperforms non-ZI shrinkage workflows and scLink.

**Figure 2:**
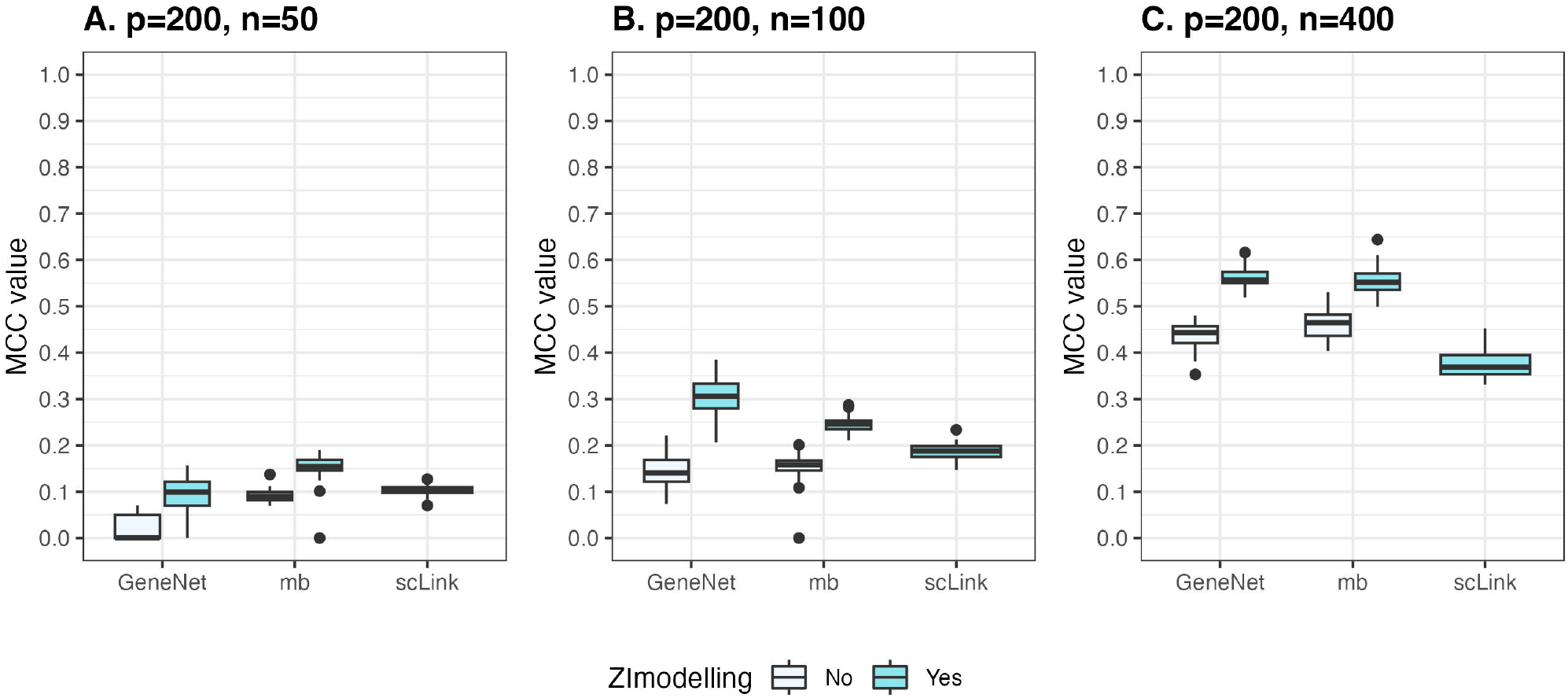
Performance of zero-inflated (ZI modelling) Stein-type and Lasso shrinkage frameworks compared to scLink workflow in zero-inflated simulated scRNAseq data (p number of features/genes, n number of observations/cells).

### 3.4 Implementation in gene network analysis of experimental scRNAseq data

To validate interaction networks estimated from the proposed zero-inflated covariance shrinkage framework, databases curating experimental evidence of gene regulation and protein interactions from budding yeasts *Saccharomyces cerevisiae* and fission yeasts *Schizosaccharomyces pombe* are used. For gene regulatory networks, YEASTRACT and Pombase are utilised for *S.cerevisiae* and *S.pombe*, respectively [37, 38]. For proteinprotein interaction networks, STRING, IntAct and BioGRID are used to analysed reported interactions of gene products. For STRING database, only interactions having evidences in experiments and databases are considered as reported interactions. Regarding experimental scRNAseq data, high-quality count data of 127 wild-type *S.cerevisiae* cells (strain BY4741) and 108 *S.pombe* cells (strain 972h-) from the Nadal-Ribelles et al. [39], and Saint et al. [40] studies are used to contruct gene regulatory networks. Genes with expression over 60% of cells in each scRNAseq data set and present in interaction networks of each database are selected. If the number of selected genes is over 200, top 200 highly variable genes (HVGs) are chosen by using modelGeneCV2 and getTopHVGs functions in scran R package [41]. In *S.cerevisiae* scRNAseq data analysis, 200 highly variable genes are shared in all databases whilst in *S.pombe* scRNAseq data analysis, HVG detection step was only executed when comparing with STRING database. Precision, or positive predictive value (PPV) is adopted as the performance metric in experimental data analysis as only a fraction of reported interactions in database are expected to be present. Details of experimental data analysis and performance evaluation metrics is in Supplementary material 9.

In the experimental data analysis, ZI and non-ZI GeneNet shrinkage workflows are compared with scLink and Pearson correlation (PearCorr) on *S.cerevisiae* and *S.pombe* data. Pearson correlation is widely used to analyse and construct gene co-expression network [42]. Pearson correlation matrix is calculated directly from count matrix using rcorr function from Hmisc R package [43]. Pearson correlation coefficients with BenjaminiHochberg adjusted p-values smaller than 0.01 are selected for final graph. In general, 200 highly variable genes present in both *S.cerevisiae* scRNAseq data and databases are applied gene network inference methods. Number of estimated edges which are reported in each database and total estimated edges are illustrated in Supplementary Table S2. Positive predictive rate is shown in Table 2. Zero-inflated GeneNet shrinkage estimates less edges than scLink and PearCorr, however, around 30% of estimated edges are reported in interaction databases compared to traditional gene network inference method, Pearson correlation (PPV *≃* 12%). Similarly, in *S.pombe* analysis, ZIGeneNet estimates edges with higher precision than other methods. For example, 1 out of 11 edges are common between ZIGeneNet estimation and PomBase database while none of 220 edges from scLink and 1 out of 169 edges from PearCorr are reported interactions.

**Table 2:** Positive predictive rates of *S.cerevisiae* scRNAseq data analysis.

| Methods | Positive predictive rates |
| --- | --- |
| ZIGeneNet | 32/108 $\simeq$ 29.63% |
| GeneNet | 10/78 $\simeq$ 12.82% |
| scLink | 46/355 $\simeq$ 12.96% |
| PearCorr | 1242/10572 $\simeq$ 11.75% |

Networks inferred by ZIGeneNet and PearCorr on *S.cerevisiae* scRNAseq data are visualised in Figure 3. Histone proteins encoded by HTA1, HTA2, HTB1, HTB2, HHT1 genes form a separate cluster in the ZIGeneNet network [44]. Hsp70 and hsp90 chaperones encoded by STI1, HSC82, SSA2, HSP82 also connect into a discrete cluster in the network [45]. Moreover, activity of TUP1 transcription factor is displayed to modulate the expression of hexokinase expressing gene EMI2 which has been reported in a comprehensive analysis of transcription factor activity in *S.cerevisiae* [46]. Compared to Pearson correlation matrix approach, partial correlation matrix estimated by zero-inflated GeneNet shrinkage framework has sparser network including activities of different protein complexes, enzymes and transcription factors.

**Figure 3:**
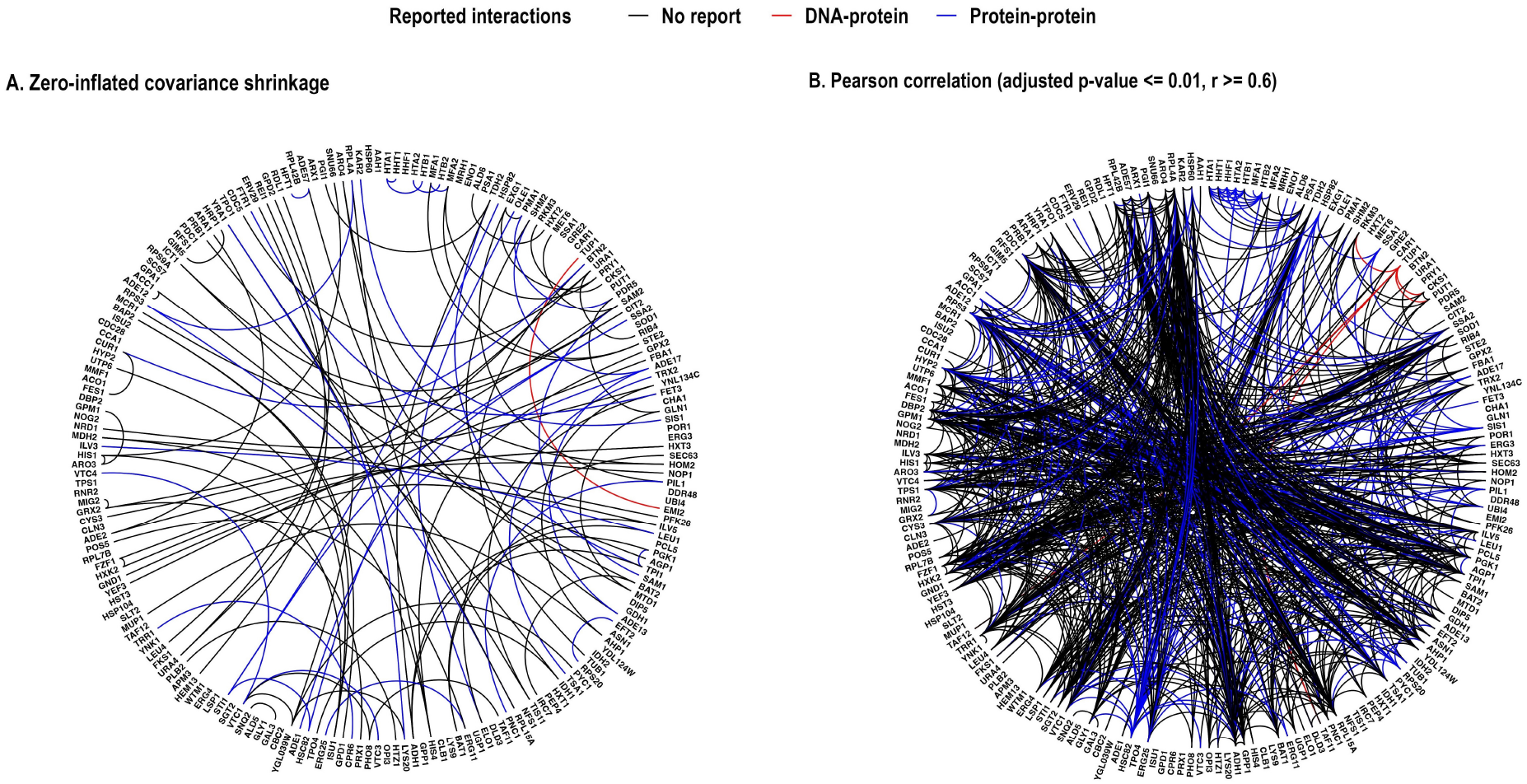
Estimated networks from zero-inflated covariance shrinkage workflow versus Pearson correlation workflow with coefficients (r) over 0.6 in analysis of *S.cerevisiae* scRNAseq data.

## 4 Conclusions

Gene regulatory network inference algorithms and allow the extraction of network information from single-cell gene expression data that can be used to further our understanding of molecular networks and potentially act as a screening step to select potential proteins for downstream analysis.

We demonstrate that our proposed approach that uses a network inference method specific to scRNAseq data gives higher performance on simulated data, and higher precision on experimental scRNAseq data than those that do not explicitly model zero-inflation. We also show that by using Stein type shrinkage and a negative binomial model of scRNAseq data that takes into account zero-inflation of counts, we can improve performance over the previously developed scLink methodology.

Due to the computational speed of Stein type shrinkage approaches, in future work we will explore the possibility of performing large numbers of local network inference tasks in scRNAseq data, to make this approach applicable to data where there are potentially continuous trajectories of cell types present, such as in developmental studies [47].

## Supplementary materials

### 1 Lasso-type shrinkage approach

In the Lasso-type shrinkage workflow, estimated adjacency matrices corresponding to each regularization parameter are produced after the shrinkage step. In terms of graph selection approaches in Lasso-type shrinkage frameworks, the STARS algorithm improves performance over the RIC algorithm when analysing in normally distributed and singlecell RNA sequencing (scRNAseq) data with mb or glasso shrinkage algorithms and no data transformation in simulation (Supplementary Figure S1 & S2). Hence, the STARS algorithm is chosen for optimal graph selection step in Lasso-type shrinkage framework[48].

**Figure S1:**
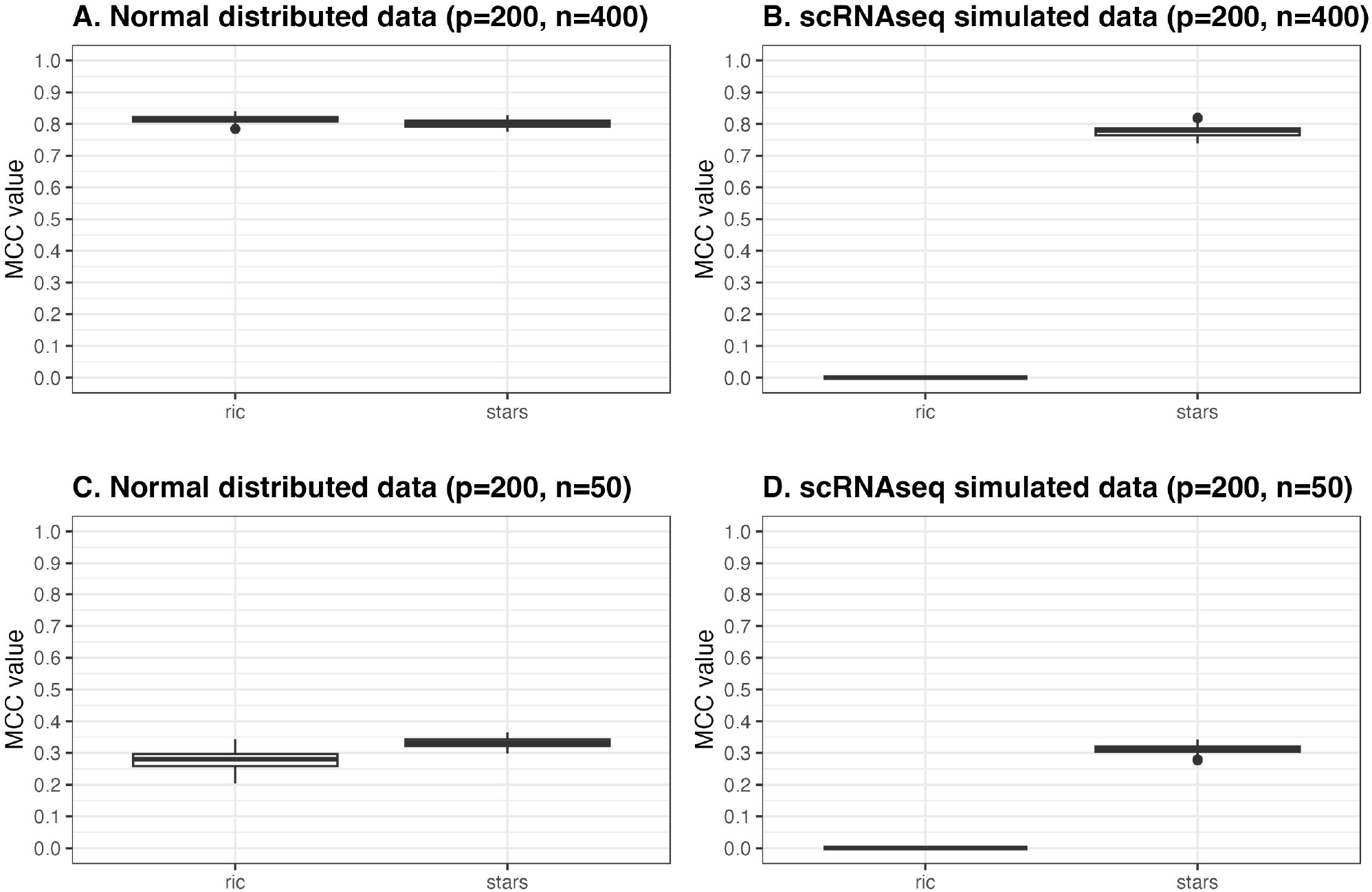
Comparison analysis between ric and stars algorithms in graph selection of mb shrinkage algorithm.

**Figure S2:**
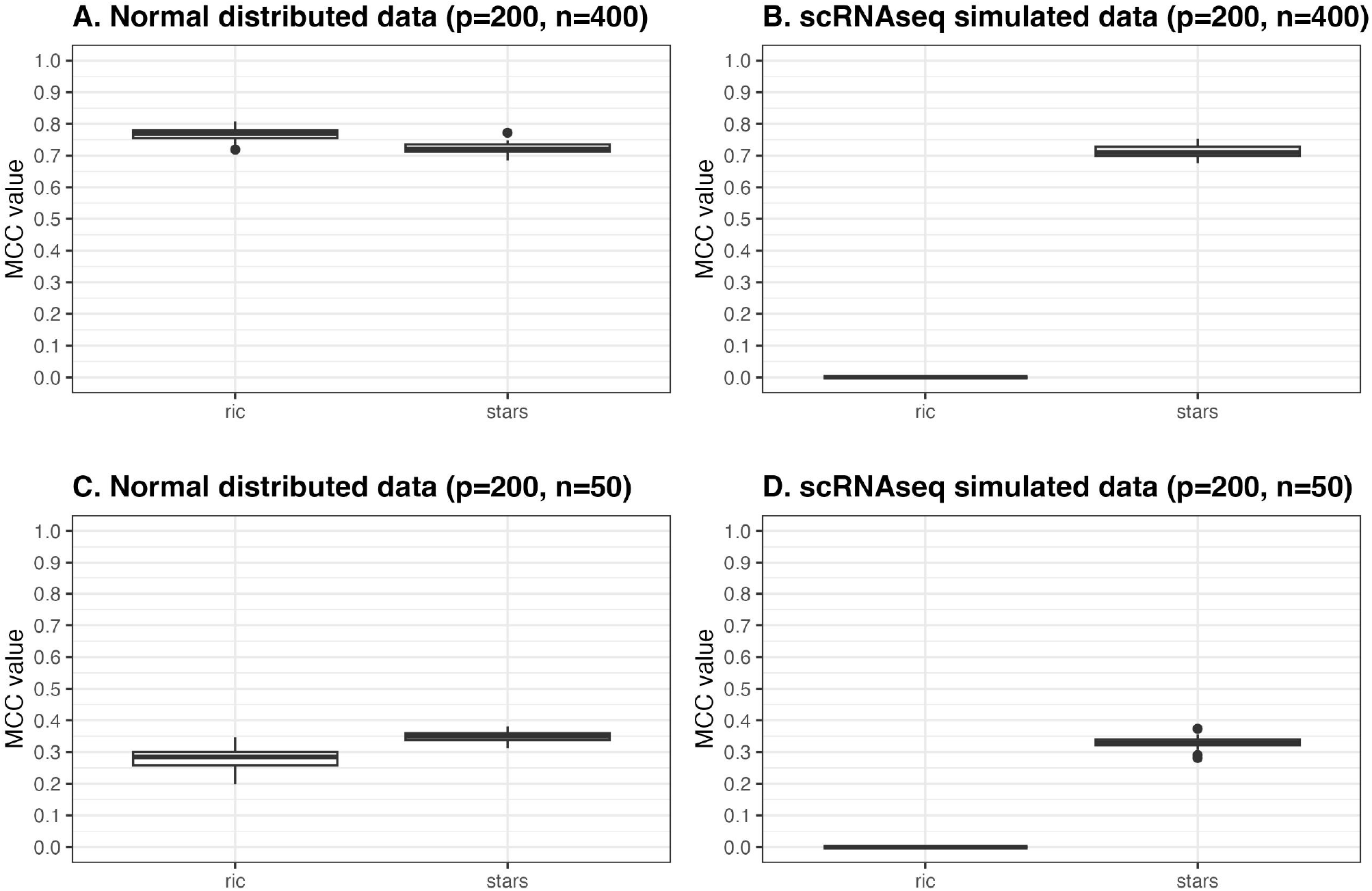
Comparison analysis between ric and stars algorithm in graph selection of glasso shrinkage algorithm.

### 2 Stein-type shrinkage approach

**Table S1:**
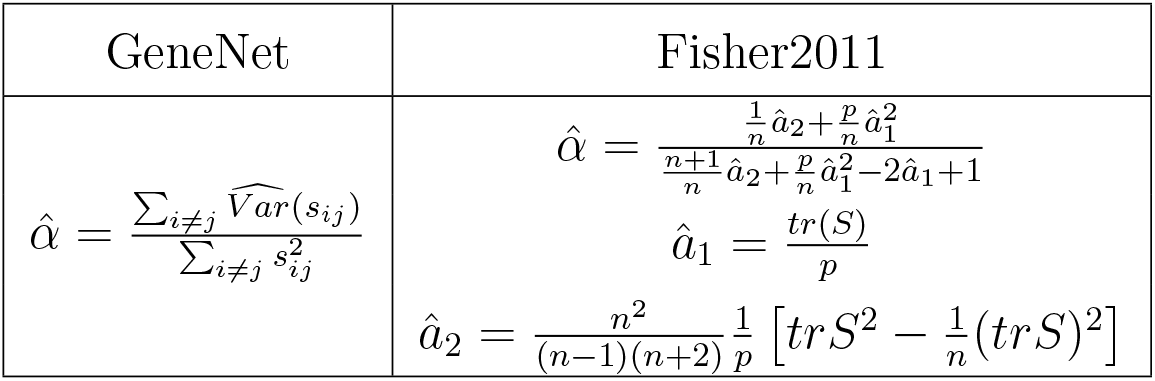
Shrinkage intensity formula of Stein-type shrinkage. Identity matrix is chosen as target matrix and optimal shrinkage intensity is derived from minimizing risk function of Frobenius loss. The shrinkage intensity formulas from Schäfer and Strimmer [16] (GeneNet) and Fisher and Sun [25] (Fisher2011) are used in the study.

| GeneNet | Fisher2011 |
| --- | --- |
| $\hat{\alpha} = \frac{\sum_{i \neq j} \widehat{Var}(s_{ij})}{\sum_{i \neq j} s_{ij}^2}$ | $\hat{\alpha} = \frac{\frac{1}{n}\hat{a}_2 + \frac{p}{n}\hat{a}_1^2}{\frac{n+1}{n}\hat{a}_2 + \frac{p}{n}\hat{a}_1^2 - 2\hat{a}_1 + 1}$ $\hat{a}_1 = \frac{tr(S)}{p}$ $\hat{a}_2 = \frac{n^2}{(n-1)(n+2)} \frac{1}{p} \left[ tr(S^2) - \frac{1}{n}(tr(S))^2 \right]$ |

### 3 Covariance matrix shrinkage workflows

**Figure S3:**
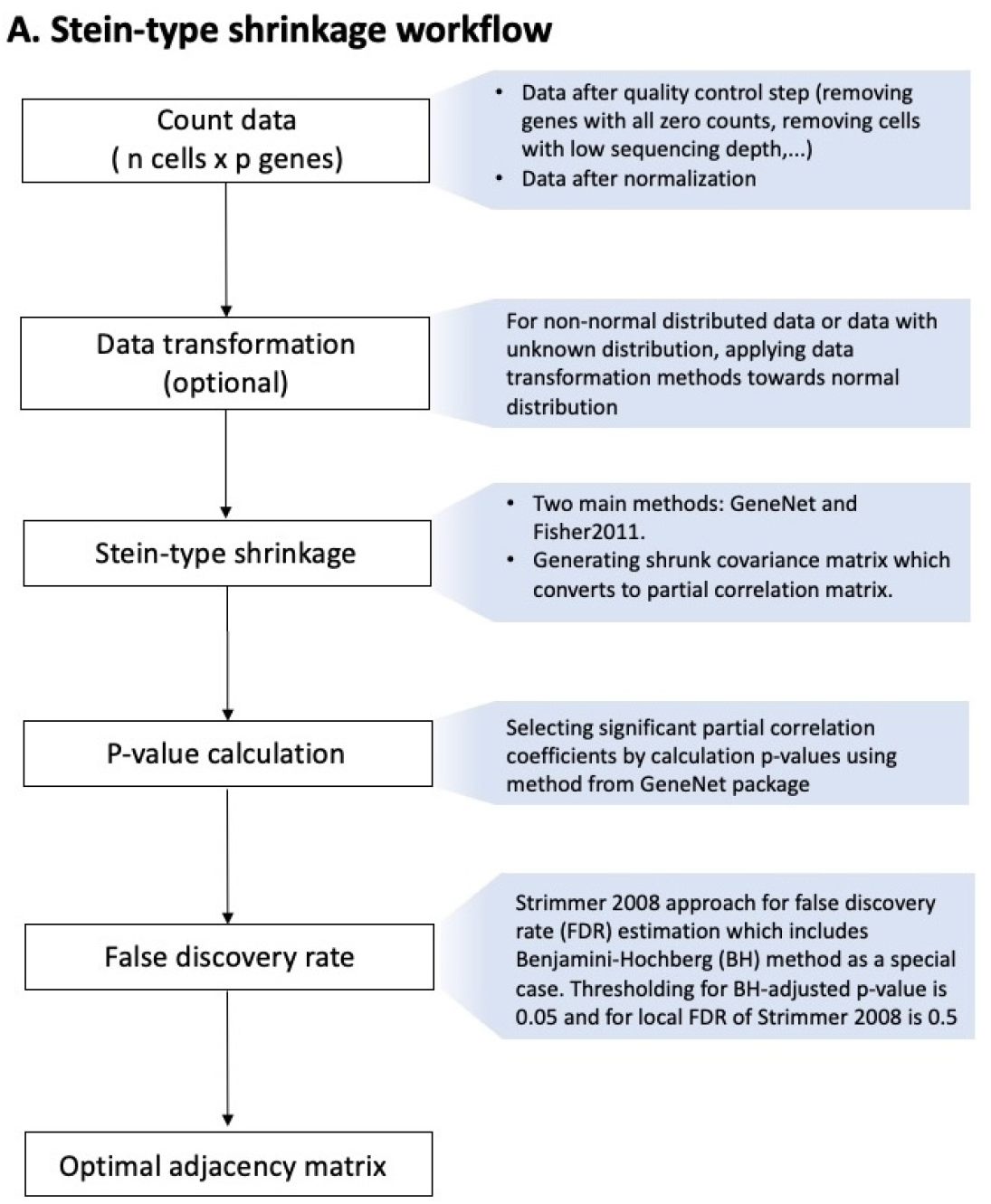
Covariance matrix shrinkage workflows of Stein-type shrinkage. Inverse covariance matrix shrinkage estimation starts with processing count data with data transformation. In Stein-type shrinkage workflow, GeneNet and Fisher2011 are implemented and partial correlation matrix is estimated from shrunk covariance matrix. Significant coefficients in partial correlation matrix are selected based on adjusted p-values and considered as edges in final adjacency matrix.

**Figure S4:**
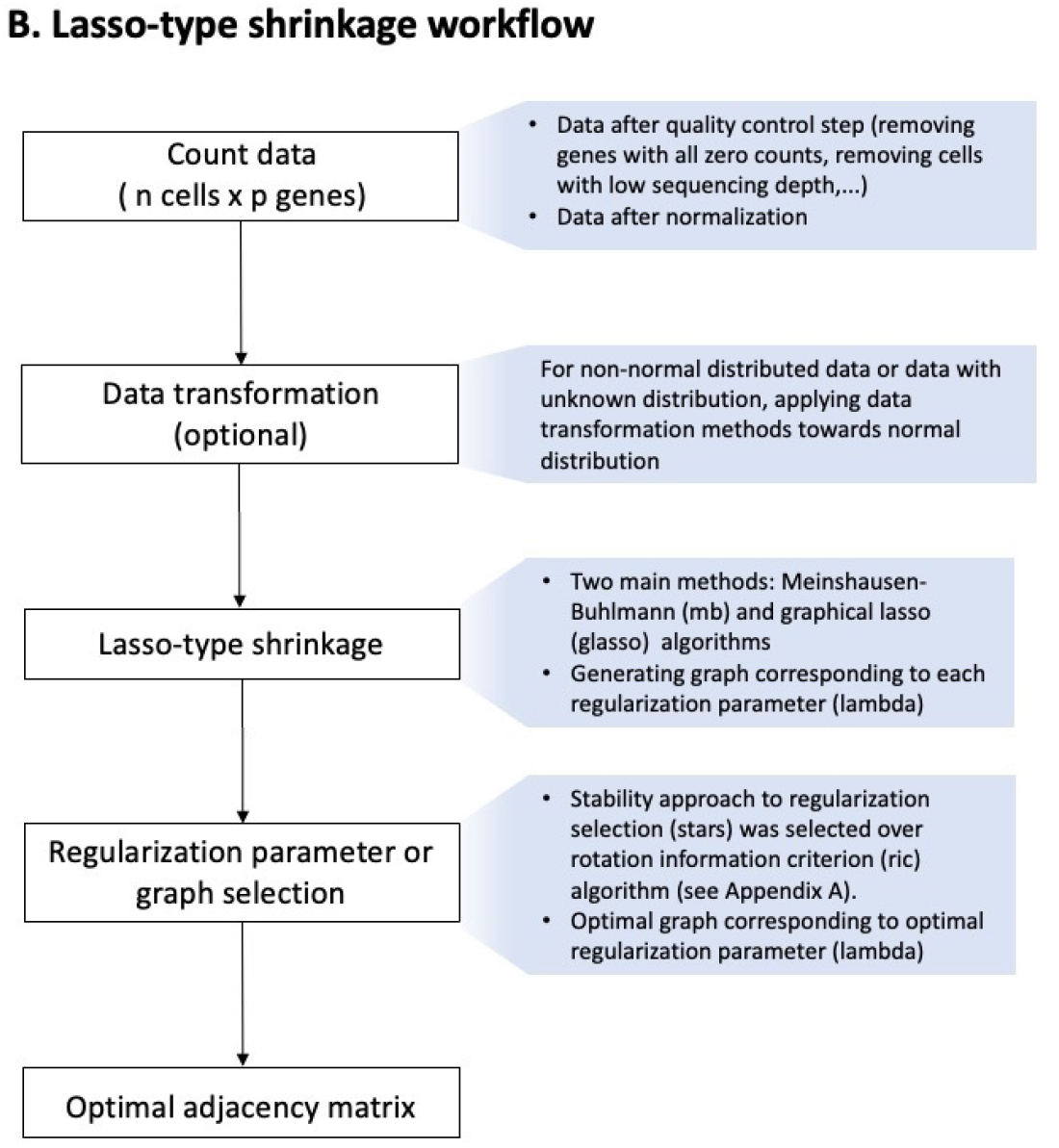
Covariance matrix shrinkage workflows of Lasso-type methods. Lasso-type shrinkage based on L1 regularization consists of Meinshausen-Buhlmann (mb) and graphical lasso (glasso) algorithms which generates different graphs corresponding to different regularization parameters. Optimal graph is selected by STARS algorithm [48]

### 4 Zero-inflated covariance matrix shrinkage workflow

To account for presence of excessive zero counts in scRNAseq data, UMI count data is transformed into zero-inflated z scores for partial correlation matrix estimation. Zeroinflated z-score calculation step can be integrated in all shrinkage workflows. Details of each step is illustrated in Supplementary Figure S5B. Importantly, at zero-inflated negative binomial modelling step, value of mean parameter (*μ*) can be the product of optimizing parameter *μ*^*′*^ and scaling factors s. Scaling factors are used in data normalization for sequencing data. They are calculated using quickCluster function for cell clustering, computeSumFactors for estimating factor value in each cluster (scran R package) [41].

**Figure S5:**
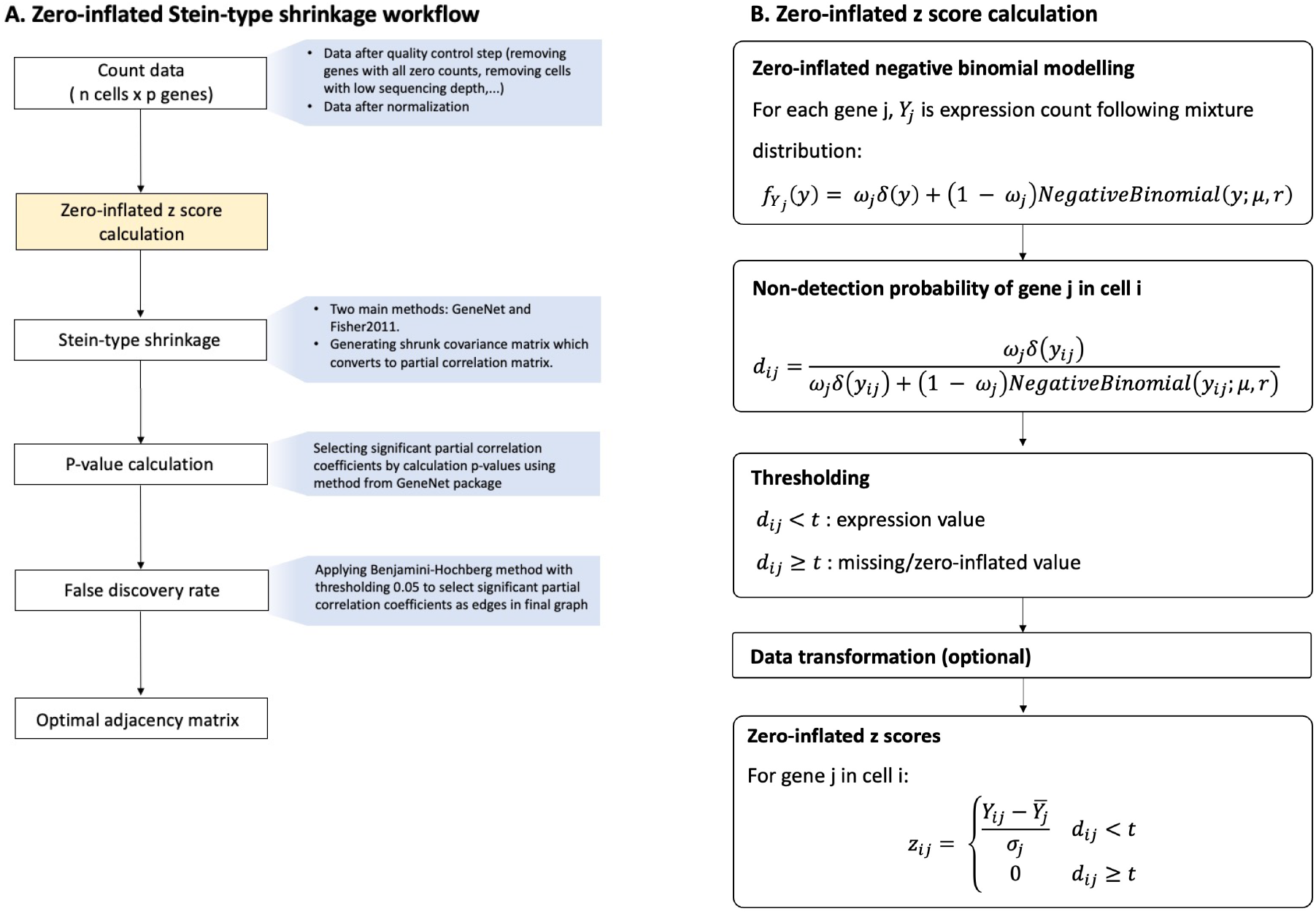
Zero-inflated covariance matrix shrinkage workflow. To adjust the presence of extra zero counts resulted from dropout events, zero-inflated z scores were calculated and used as an input for Stein-type shrinkage. Zero-inflated negative binomial is fitted to stratify zero counts from dropout event and “biological” zero counts. After the fitting, non-detection rate (*d*_*ij*_) is estimated in each cell for each gene. Thresholding (default t=0.5) is applied in which counts with *d*_*ij*_ *≥t* are considered missing or zero-inflated values. Z scores are calculated for all counts and scores of zero-inflated values are set to 0.

### 5 Simulation of scRNAseq data

**Figure S6:**
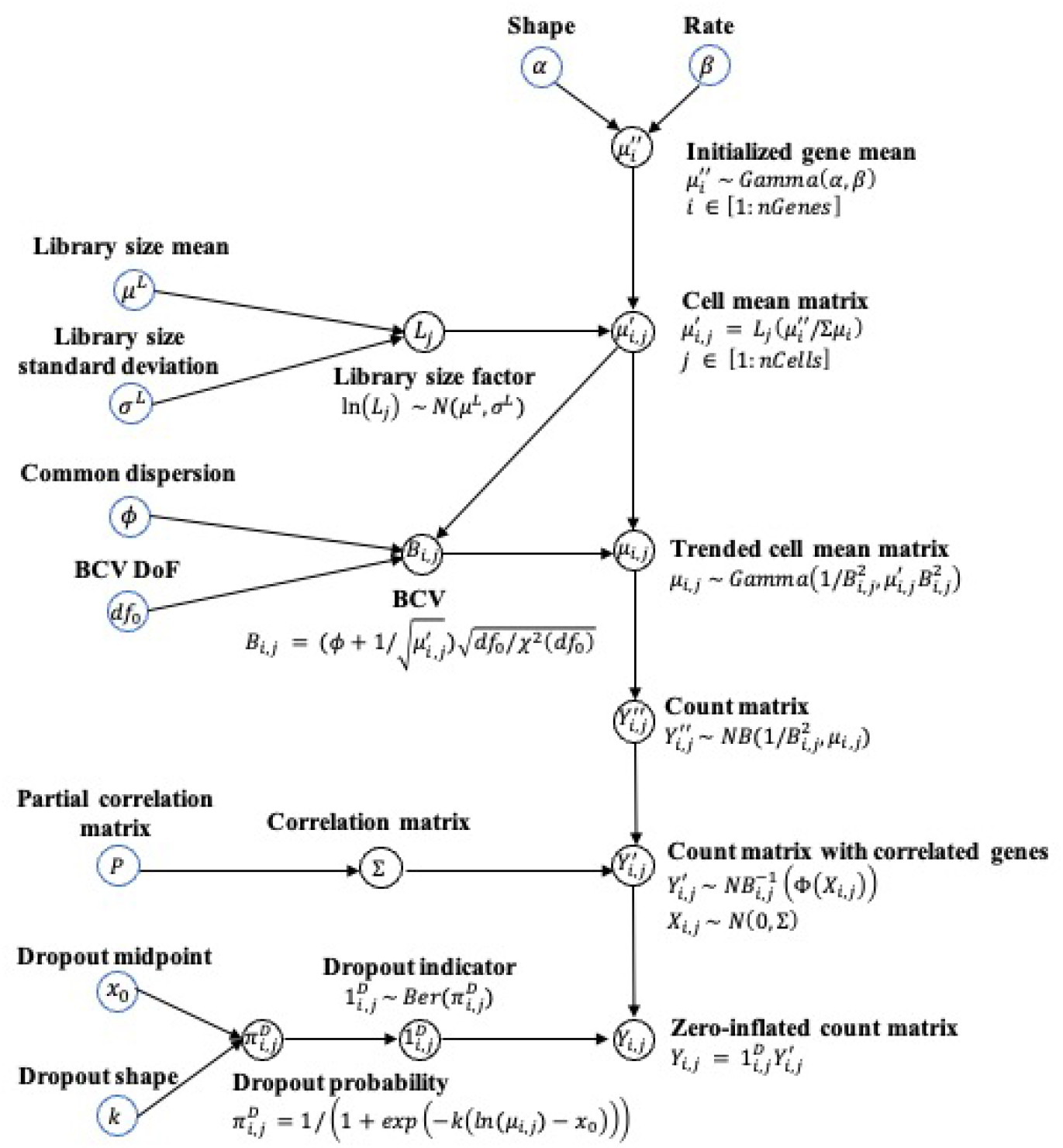
Simulation model. Blue circles represent input parameters whilst black circles indicate calculating or simulating steps. Simulation model is based on models from splatter and ESCO R packages [32, 49]. Gene mean is simulated from Gamma distribution. Means of gene expression are converted into matrix in which library size and Biological Coefficient of Variation (BCV) are used to adjust the expression in each cell. Final mean matrix is used to simulate count matrix from negative binomial distribution. Input partial correlation matrix is converted to correlation. Next, co-expression gene network is embedded in count matrix by using copula and dropout event is introduced through logistic expression depending on gene mean.

### 6 Data transformation and p-value estimation

**Figure S7:**
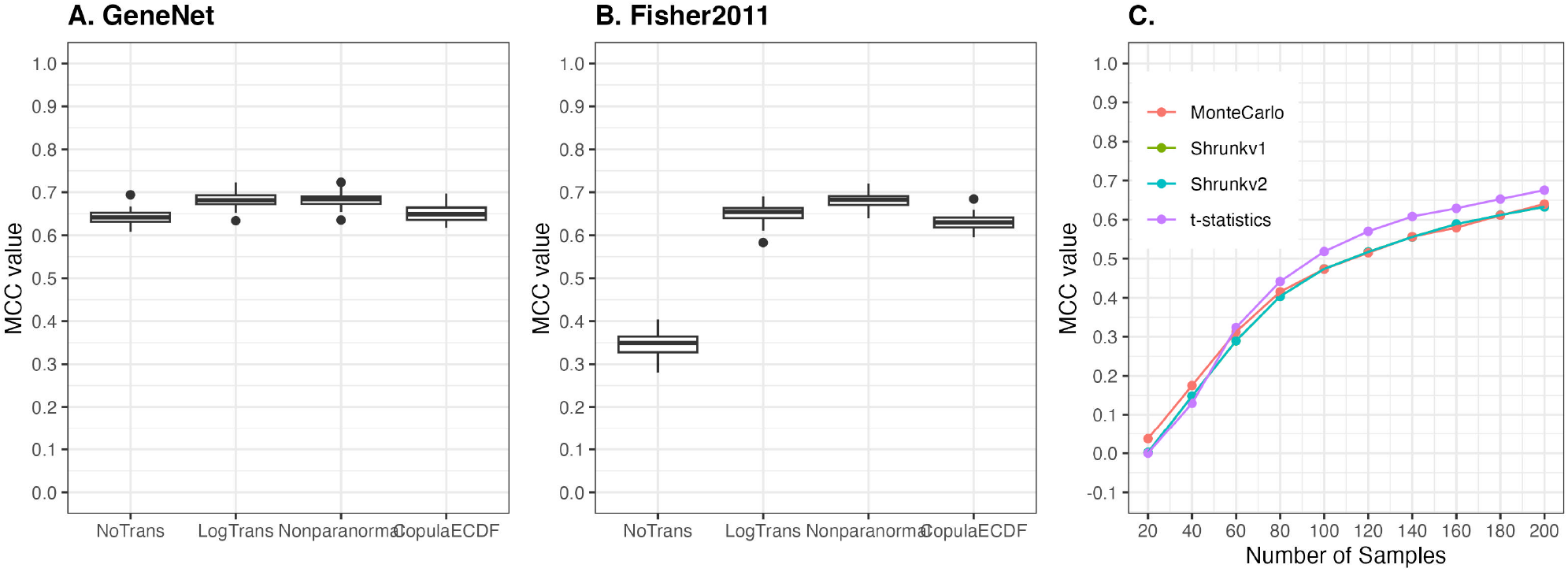
Comparative analysis of different approaches in data transformation and p-value calculations on partial correlation matrix. A & B, Three transformation methods were applied on simulated scRNAseq data with p = 200, n = 200, sequencing depth = 50000 reads and partial correlation matrix was estimated by Fisher2011 algorithm. Compared to no transformation (NoTrans), performance of Fisher2011 was improved most by nonparanormal approach (Nonparanormal) from [29] and log transformation (LogTrans). C, Different models to calculate p-value from shrunk partial correlation matrix were applied on simulated data (p = 200, sequencing depth = 50000 reads, GeneNet shrinkage). Models used were t-statistics, Monte Carlo method (MonteCarlo), shrunk probability density models for shrunk partial correlation matrices (shrunkv1 and shrunkv2 from [19] and [31], respectively).

### 7 Zero proportion in experimental scRNAseq data

**Figure S8:**
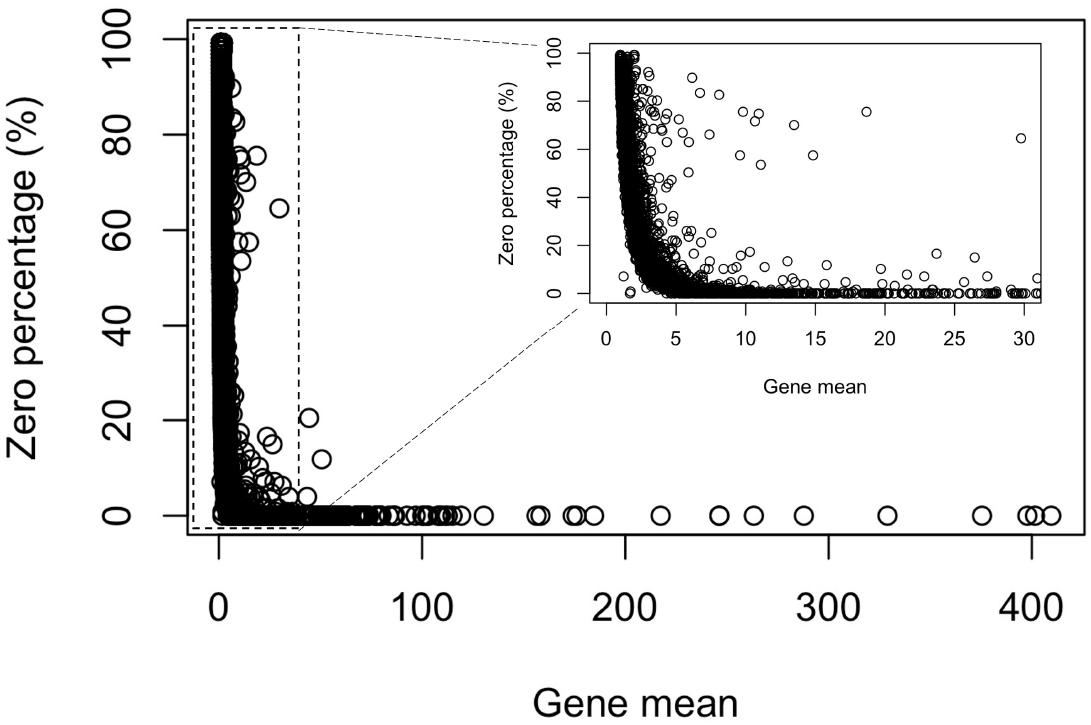
Zero proportion in *S.cerevisiae* scRNAseq data. Accession number: GSE122392 [39].

**Figure S9:**
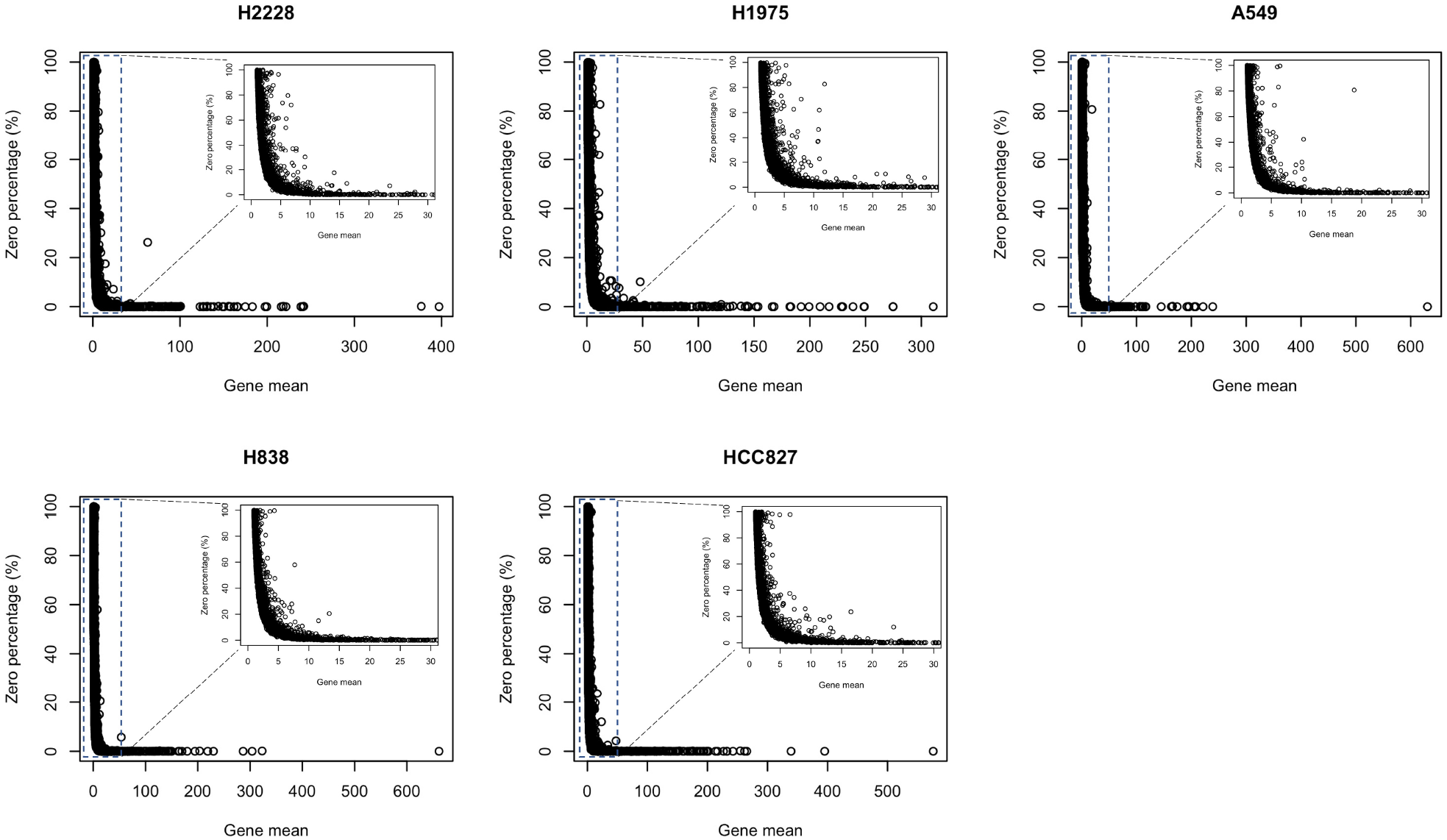
Zero proportion in benchmark datasets (10x Genomics protocol). Accession number: GSE118767 [**tian2019benchmarking**].

### 8 Performance of zero-inflated modelling in simulated scRNAseq data

**Figure S10:**
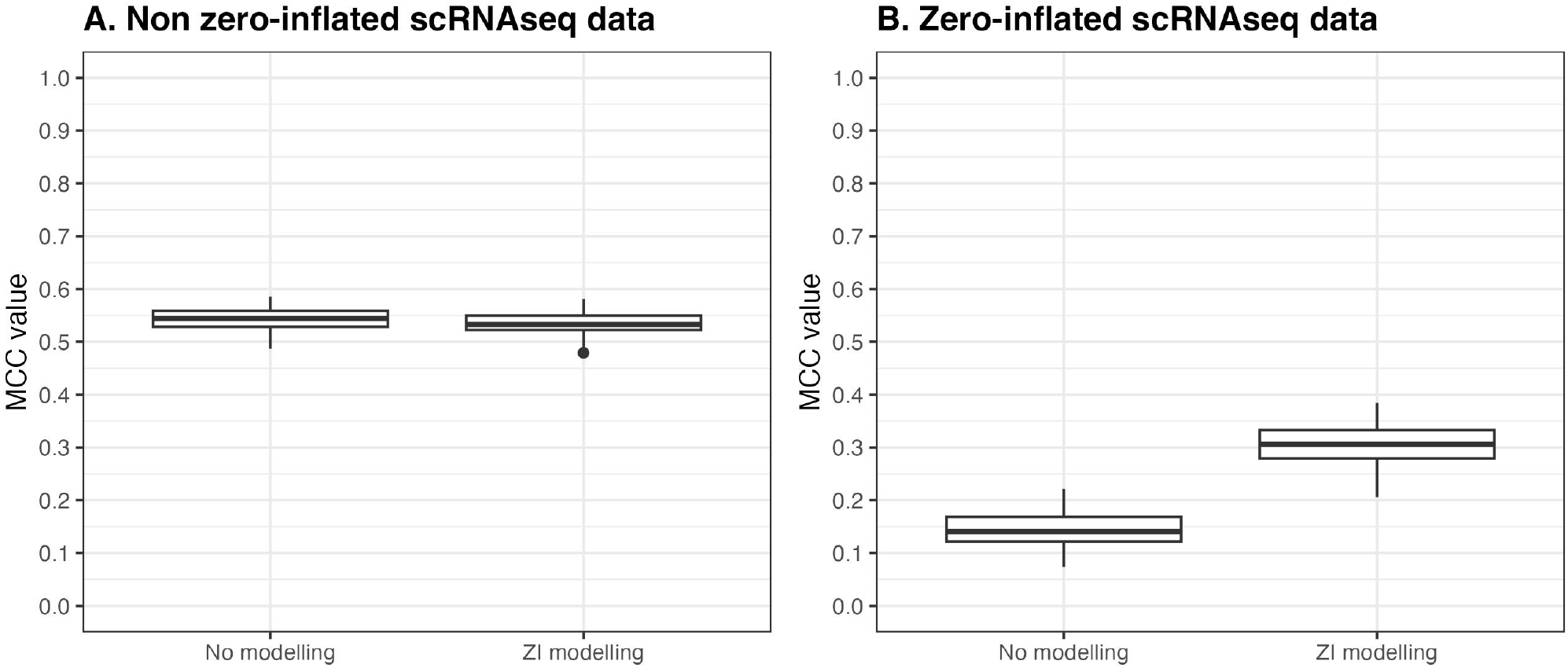
Performance with and without zero-inflated modelling in Stein-type shrinkage workflow in non-zero inflated (A) and zero-inflated (B) simulated scRNAseq data. (GeneNet, p=200, n=100)

**Figure S11:**
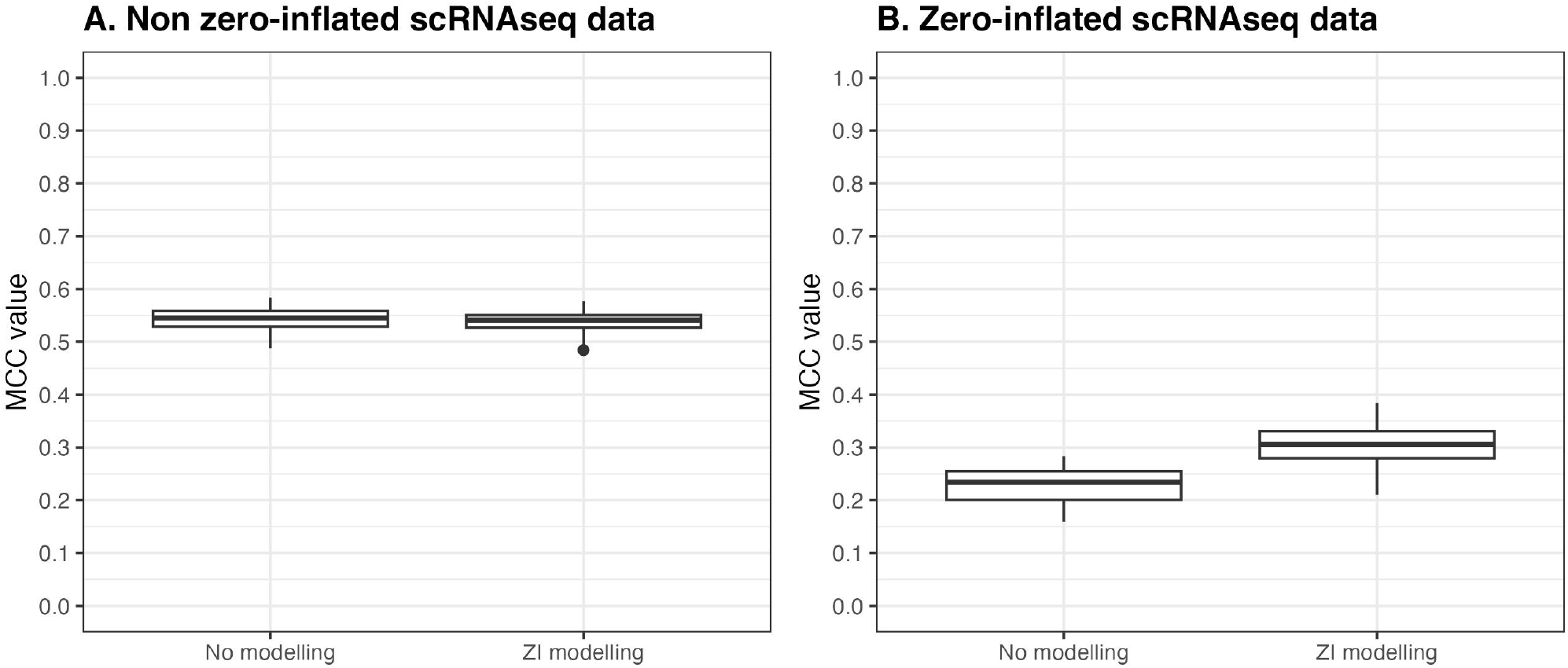
Performance with and without zero-inflated modelling in Stein-type shrinkage workflow in non-zero inflated (A) and zero-inflated (B) simulated scRNAseq data. (Fisher2011, p=200, n=100)

### 9 Experimental scRNAseq data analysis

In covariance or correlation matrix calculation, observations are assumed to be independent and identically distributed (iid) and in case of scRNAseq data, observations are cells [50]. To avoid the violation of iid assumption, covariance or correlation matrix should be calculated on data of cells from the same population, growing under the same condition [51]. In *S.cerevisiae* scRNAseq data, 127 cells of same strain BY4741 were grown in same condition and included in network analysis (accession number GSE122392) [39]. In *S.pombe* data, 108 972hcells from dataset 1712 1 were grown and maintained in same condition are chosen (accession number E-MTAB-6825) [40].

### Performance evaluation metrics

For evaluation of the network inference schemes on simulated data, the Matthews correlation coefficient (MCC) is used, which is a contingency matrix method to calculate the Pearson product-moment correlation coefficient between prediction and reference values [52]. The MCC score is calculated as follows,

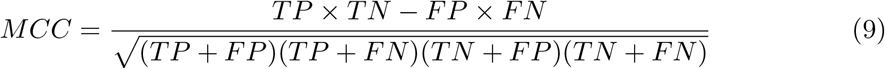

where TP is the count of true positive predictions, TN is the count of true negative predictions, FP represents the count of false positives, and FN the number of false negative predictions. The MCC ranges from *–*1 to 1, which indicates the worst and perfect classification respectively. The MCC is commonly applied in binary classification evaluation for imbalanced data [52]. As gene regulatory networks are sparse relative to the total possible number of edges, the task of gene regulatory network inference has a large class imbalance, with the negative class (non-edges) being much larger than the positive (edges).

Gene regulatory and protein-protein interaction database combine results of organisms living different experimental conditions. Therefore, not all of reported interactions in these databases are likely to be present in one experimental condition. Hence, in experimental data analysis, to compare estimated interactions to reported interactions in databases, fraction of reported interactions in all estimated edges is central focus. Precision or positive predictive value (PPV) is adopted and calculated as follows:

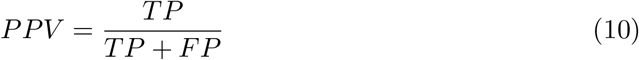

**Table S2:** Comparison of experimental results with gene network databases (number of reported interactions/number of total estimated edges).

|  | Gene regulatory<br>networks | Protein-protein networks |  |  |
| --- | --- | --- | --- | --- |
|  |  | STRING | IntAct | BioGRID |
| *S.cerevisiae* |  |  |  |  |
| Number of genes | 200 | 200 | 200 | 200 |
| ZIGenNet | 1/108 | 20/108 | 17/108 | 24/108 |
| GeneNet | 0/78 | 6/78 | 8/78 | 8/78 |
| scLink | 9/355 | 24/355 | 22/355 | 27/355 |
| PearCorr | 27/10572 | 146/10572 | 515/10572 | 803/10572 |
| *S.pombe* |  |  |  |  |
| Number of genes | 77 | 200 | 194 | 166 |
| ZIGeneNet | 1/11 | 74/303 | 2/213 | 3/288 |
| GeneNet | 1/48 | 24/89 | 2/270 | 2/335 |
| scLink | 0/220 | 13/303 | 0/22 | 2/185 |
| PearCorr | 1/169 | 87/247 | 0/795 | 1/424 |

